# Microtubule association of TRIM3 revealed by differential extraction proteomics

**DOI:** 10.1101/2023.07.27.549915

**Authors:** Hannah L. Glover, Marta Mendes, Joana Gomes-Neto, Emma V. Rusilowicz-Jones, Daniel J. Rigden, Gunnar Dittmar, Sylvie Urbé, Michael J. Clague

## Abstract

The microtubule network is formed from polymerised Tubulin subunits and associating proteins, which govern microtubule dynamics and a diverse array of functions. To identify novel microtubule binding proteins, we have developed an unbiased biochemical assay, which relies on the selective extraction of cytosolic proteins from cells, whilst leaving behind the microtubule network. Candidate proteins are linked to microtubules by their sensitivities to the depolymerising drug Nocodazole or the microtubule stabilising drug, Taxol, which is quantitated by mass spectrometry. Our approach is benchmarked by co-segregation of Tubulin and previously established microtubule-binding proteins. We then identify several novel candidate microtubule binding proteins, from which we have selected the ubiquitin E3 ligase TRIM3 (Tripartite motif-containing protein 3) for further characterisation. We map TRIM3 microtubule binding to its C-terminal NHL-repeat region. We show that TRIM3 is required for the accumulation of acetylated Tubulin, following treatment with Taxol. Furthermore, loss of TRIM3, partially recapitulates the reduction in Nocodazole-resistant microtubules characteristic of Alpha-Tubulin Acetyltransferase 1 (ATAT1) depletion. These results can be explained by a decrease in ATAT1 following depletion of TRIM3 that is independent of transcription.

## Introduction

Microtubules (MTs) are formed from α-and β-Tubulin heterodimer subunits which polymerise to form hollow fibres, making up one of three major cytoskeletal elements (Wickstead and Gull, 2011). They are polar structures, defined by a growing plus-end oriented towards the cell periphery and a minus-end which is usually stabilised at the microtubule organising centre (MTOC) (Mitchison, 1993). Microtubule networks are able to grow and shrink via continuous rounds of polymerisation and depolymerisation, allowing for constant cytoskeleton remodelling derived from their dynamic instability (Mitchison and Kirschner, 1984). Consequently, they contribute to a wide variety of cellular functions such as maintaining cell shape, cell movement, cell division and provide the main structural unit for flagella and cilia (Nogales, 2000). They also provide tracks, along which motor proteins can travel, to distribute their cargo such as organelles and membranous vesicles (Barlan and Gelfand, 2017; Bloom and Endow, 1995).

Numerous microtubule associated proteins (MAPs) account for this diverse array of functions. Some specifically bind to the end of microtubules to either control their attachment to cellular structures (-end) or their dynamics (+end). The first MAPs were discovered during the development of protocols for Tubulin purification (Sloboda et al., 1975; Souphron et al., 2019). Similarly, the protein Tau was identified from porcine brain extracts, as a factor governing microtubule polymerisation (Weingarten et al., 1975). Modern LC-MS/MS techniques provide a large scale, global and unbiased method for identifying MAPs. Prior MAP-centric proteomic studies have been performed using a variety of species. Over 250 MAPs were identified from early Drosophila embryos using Taxol-and GTP-stabilised endogenous Tubulin preparations, followed by 2D gel electrophoresis (Hughes et al., 2008). In a separate study, macrophage extracts were incubated with the microtubule stabilising drug Taxol and purified bovine brain Tubulin, with candidate MAPs being identified by spectral counting (Patel et al., 2009). In 2010, more than 300 proteins from meiotic *Xenopus* egg extracts were found to bind to Taxol-stabilised bovine brain Tubulin (Gache et al., 2010). A final example identified over 1000 proteins binding *in vitro* to the metaphase spindle isolated from CHO cells, using Multi-dimensional Protein Identification Technology (Bonner et al., 2011).

The majority of these studies preceded major advances in the sensitivity of mass spectrometry instruments and the adoption of isotopic labelling procedures (e.g. SILAC, DML, TMT) that provide more quantitative data. Furthermore, many involve the manipulation of the microtubule network *in vitro* and/or the addition of exogenous Tubulin, thereby incompletely capturing the intracellular architecture and environment. In this study, we take a different approach, by identifying the proteins that remain bound to a residual microtubule network, assembled under physiological conditions, following the extraction of cytosol. Our principal criterion for presuming microtubule association is a sensitivity to the depolymerising drug Nocodazole. We describe the development of this methodology and the general features of our results. Our analysis revealed several novel candidate microtubule binding proteins including the ubiquitin E3-ligase, TRIM3 (Tripartite motif-containing protein 3), for which we present a detailed characterisation of its association and describe effects on microtubule properties.

## Results

Nocodazole interferes with microtubule polymerisation, leading to a rapid loss of the microtubule network upon application to cells in culture (De Brabander et al., 1977). Conversely, Taxol induces the assembly and stabilisation of microtubules (De Brabander et al., 1981). Following extraction of the cytosol from cells, leaving behind the microtubule network, proteins which show sensitivity to either of these drugs become strong candidates for microtubule association. By labelling each condition with amino acids bearing different stable isotopes, residual non-microtubule related (or drug insensitive) proteins are simply discounted by virtue of their 1:1 ratios, which are derived from subsequent mass spectrometry analysis. We set out to determine this form of MAPome which in distinction to previous approaches reflects the microtubule status in living cells.

### Optimisation of the microtubule extraction protocol

Our initial approach involved an iteration of a previously described method to extract microtubule-associated proteins directly from U2OS cells (Duerr et al., 1981). A buffer which lyses cells, whilst maintaining microtubules (Lysis and Microtubule Stabilisation buffer, LMS), is added to remove cytosolic proteins and small molecules, including any free Tubulin present. Vehicle-treated cells are depleted of cytosol, whereas the intact microtubules and MAPs are retained. These can then be visualised by immunofluorescence microscopy or immunoblotting. For comparison, we used a condition pre-treated with the microtubule depolymerising drug Nocodazole, which provides us with a microtubule-depleted residual fraction. We first optimised the Nocodazole incubation conditions required to achieve maximum microtubule depolymerisation. U2OS cells were treated with increasing concentrations of up to 12 µM for varying times at 37°C (Figure 1A, B). On this basis we selected a concentration of 6 µM for one hour as the optimal treatment although a minor fraction of microtubules remained resistant to depolymerisation. Rather than use high calcium conditions to release MAPs after cytosol extraction as described by Duerr et al. (Duerr et al., 1981), we have simply recovered the cell remnants (± prior Nocodazole extraction) with 8 M urea lysis buffer. Prior to a large scale mass spectrometry analysis, we checked that the established MAPs, EML4 and MAP4, demonstrated the expected behaviour by Western blotting and immunofluorescence respectively (Figure 1C, D). A schematic of the fully optimised protocol is shown in Figure 1E.

**Figure 1.**
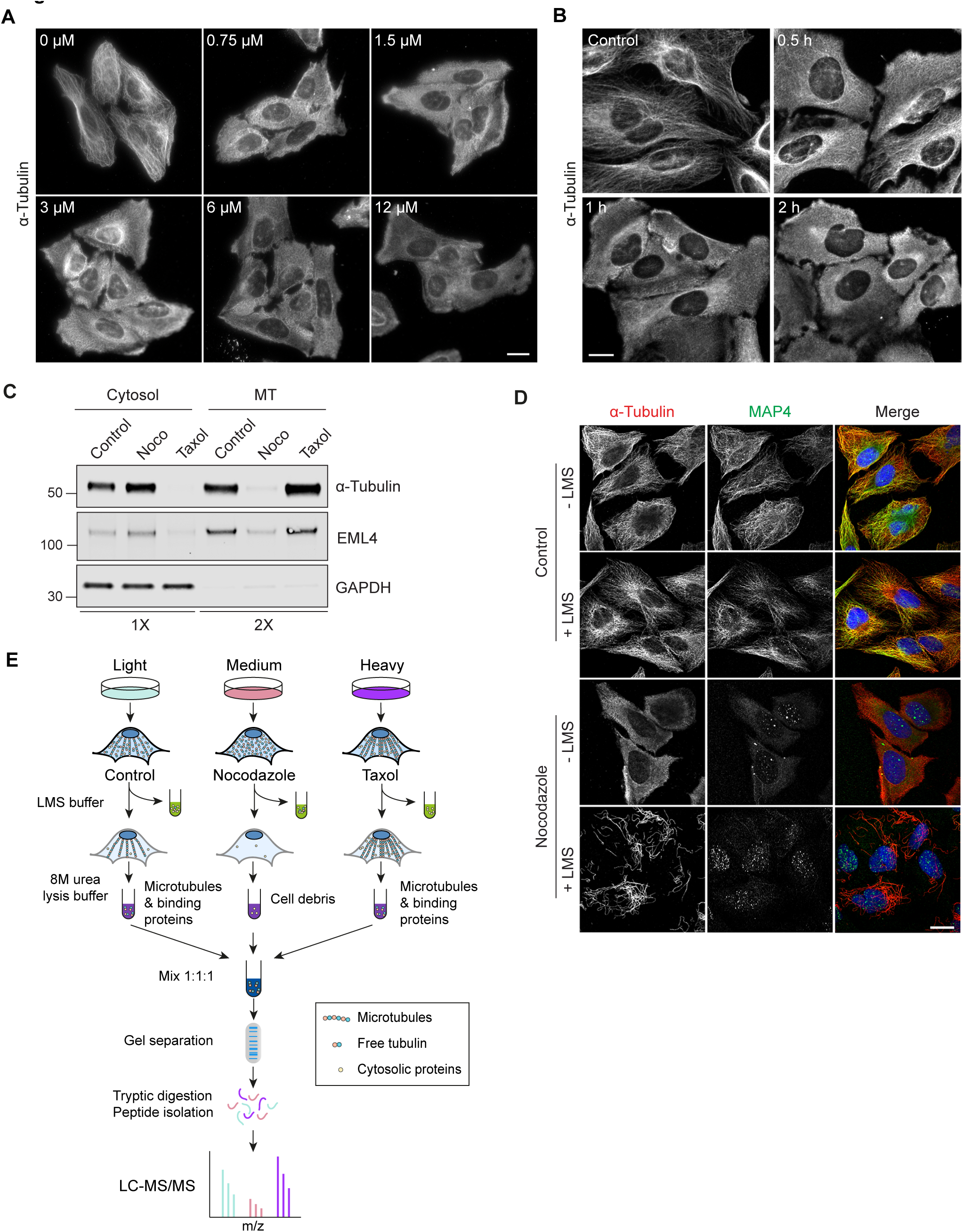
Establishing the MAPome identification work flow. **A:** U2OS cells were treated with indicated concentrations of Nocodazole at 37°C for 30 min. Cells were fixed with methanol and stained for α-Tubulin. Scale bar: 20 µm. Images were obtained using a Nikon Ti Eclipse microscope, with a CFI Plan Apo 40x objective. **B**: U2OS cells were treated with 6 μM Nocodazole for 0.5, 1 or 2 h. Cells were fixed with methanol and stained for α-Tubulin. Scale bar = 10 μm. Images were obtained using a Nikon Ti Eclipse microscope, with a CFI Plan Apo 60x objective. **C:** U2OS cells were treated with vehicle (DMSO), Nocodazole (6 µM, 1 h) or Taxol (6 µM, 30 min). Lysis and microtubule stabilisation buffer (LMS) was added (5 min, 4°C) to remove cytosolic proteins and residual cellular material (MT;microtubules) was then collected following 8 M urea lysis buffer extraction. Samples were analysed by western blot as indicated, loading twice as much MT-fraction as Cytosol). **D:** U2OS cells were treated with vehicle (DMSO) or Nocodazole for 1 h. LMS buffer was then added for 5 min at 4°C. Cells were either fixed immediately or first treated with LMS to extract cytosolic proteins prior to fixation in methanol and staining for α-Tubulin (red), MAP4 (green) and DAPI (blue). Scale bar: 20 µm. Images were obtained using a 3i spinning disc confocal microscope with a Plan-Apochromat 63x/1.4NA Oil Objective M27. **E:** Schematic showing the workflow for SILAC mass spectrometry analysis. U2OS cells incubated with light (Lys0, Arg0), medium (Lys4, Arg6) or heavy (Lys8, Arg10) amino acids were treated with either DMSO alone (control), Nocodazole or Taxol respectively as described in C. Cells were then treated with LMS buffer at 4°C for 5 min to remove cytosolic proteins and leave behind microtubules and their associated proteins. 8 M urea lysis buffer was then added to collect the residual cellular material including microtubules. These fractions were mixed at a 1:1:1 ratio and analysed by mass spectrometry.

### Determination of the microtubule proteome

To determine the microtubule associated proteome via quantitative mass spectrometry, we generated extracts from SILAC (Stable Isotope Labelling by Amino acids in Cell culture) labelled U2OS cells subjected to 3 different scenarios, untreated control or prior treatment with Nocodazole and Taxol respectively (Figure 1E). The 3 residual fractions were combined in a 1:1:1 ratio and analysed by mass spectrometry, for which full results are summarised in Supplementary Table 1. The log_2_ transformed ratio of Nocodazole/Control values were ranked from the highest negative score to the lowest. To confirm microtubule protein enrichment and define a cut-off point for further consideration, hits were analysed in sequential groups of 10 (1-10, 2-11….) for their enrichment score for “microtubule/microtubule-binding/microtubule cytoskeleton” using DAVID Bioinformatics Resource 6.8 until no enrichment was observed for 10 consecutive groups (Figure 2A) (Huang da et al., 2009). In effect this gives us an inferred Bayesian approach to determine a cut-off value for selecting MAP candidates from our data, for which data points of interest are indicated in Figure 2B. The distribution of the normalised Nocodazole/Control ratio (x-axis) versus the normalised Taxol/Control ratio (y-axis) is shown in Figures 2B-D. Proteins represented by data points in the top left quadrant of this graph, will have been lost due to Nocodazole treatment and preferentially retained following Taxol treatment. The plotted ratios derive from averaging across the identified constituent peptides of any given protein. The behaviour of each individual peptide from selected proteins is represented in Figure S1A. Figure S1B shows a heat map of outlying Nocodazole sensitive proteins across 3 independent repeats.

**Figure 2.**
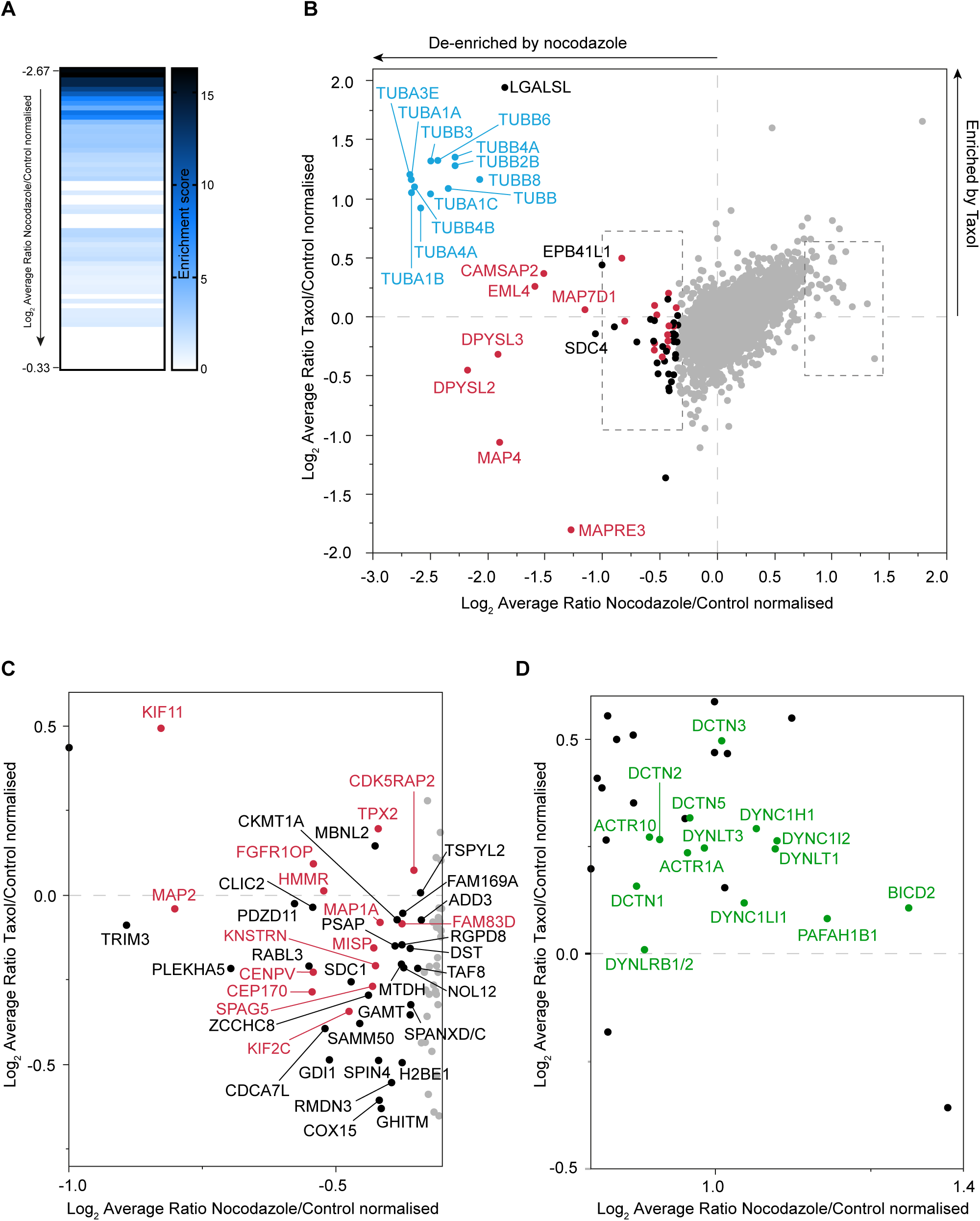
Global overview of Nocodazole and Taxol sensitivity. **A:** Heat map showing the enrichment score of microtubule proteins. Proteins ranked according to their degree of de-enrichment after Nocodazole treatment (i.e. from highest -ve X-axis value) were analysed using the functional annotation clustering tool in DAVID 6.8 (Huang da et al., 2009). Consecutive groups starting with proteins ranked 1-10 followed by hits 2-11 and so on, were assessed for their enrichment score for microtubule association. This analysis was performed in iterative fashion until no enrichment was evident for >10 consecutive groups. **B:** The log_2_ average normalised ratio of all extracted proteins in Control (DMSO-only treated; light) cells compared to Nocodazole (medium, x-axis) and Taxol (heavy, y-axis) treatments. The proteins that are de-enriched in the Nocodazole fraction were defined using DAVID as in A. Tubulin proteins in blue, known microtubule binding proteins in red, other identified binding proteins in black. **C and D:** Expanded views of proteins enriched following Nocodazole treatment with known microtubule binding proteins shown in red (C) or green (D).

As expected, the strongest outliers in terms of Nocodazole sensitivity are 12 isotypes of Tubulin (labelled in blue, Figure 2B). These are also modestly enriched by Taxol treatment. We were surprised to find Galectin-related protein (LGALSL), to be so strongly aligned with this cluster. This small protein is a candidate amyotrophic lateral sclerosis (ALS) gene, which contains a Galectin domain, although most of the residues critical for carbohydrate binding are not conserved (Gelfman et al., 2019). We established that LGALSL is a *bone fide* MAP through immunofluorescence visualisation. Although there is a majority cytosolic background stain, concentration at microtubules can clearly be discerned (Figure S2A, top row, inset). This becomes even more obvious during cytokinesis, where microtubules are bundled either side of the midbody (Figure S2B). Although masked by the large cytosolic pool in control cells, the microtubule association becomes more discernible upon cytosol extraction with LMS and is disrupted by Nocodazole treatment (Figure S2A, second and third rows). Other Nocodazole sensitive outliers include EML4 (Echinoderm microtubule-associated protein-like 4), and the tightly clustered DPYSL2 and DPYSL3 (Dihydropyriminidase like 2 and 3), otherwise known as Collapsin response mediator protein-2 and -4 respectively (Figure 2B). DPYSL2 and 3 can form heterotetramers which have been shown to enable microtubule growth (Fukata et al., 2002; Tan et al., 2015). MT minus-ends that are not attached to the centrosome are frequently tethered to the Golgi apparatus and can be stabilised by calmodulin-regulated spectrin-associated 2 (CAMSAP2, sensitive to Nocodazole only, Figure 2B) (Hendershott and Vale, 2014). Amongst these proteins de-enriched by Nocodazole treatment and clearly separated from the main cloud of data-points, we find the ubiquitin E3 ligase family member Tripartite Motif Containing 3 (TRIM3), for which we provide a detailed characterisation below (Figure 2C).

Within the bottom left quadrant of Figure 2B, we identify proteins that are lost upon both Nocodazole and Taxol treatments i.e. those that associate with dynamically unstable microtubules. Prominent here is MAP4 which is displaced by Taxol treatment (Xiao et al., 2012), but also MAPRE3 (Microtubule Associated Protein RP/EB Family Member 3) otherwise known as EB3. It is a plus-end tracking protein (+TIP) that binds to the plus-end of microtubules and regulates the dynamics of the microtubule cytoskeleton.

We were surprised to find a coherent group of proteins separated from the cloud on the right hand side of the X-axis i.e. enriched by Nocodazole treatment (labelled in green in Figure 2D). This group contains multiple subunits of the cytoplasmic dynein and dynactin complexes: the minus-end directed microtubule motor protein and its essential cofactor respectively (Trokter et al., 2012; Urnavicius et al., 2015). Bicaudal D2 (BICD2) is an adaptor protein, which recruits dynein and stabilises the interaction between dynein and dynactin. Platelet Activating Factor Acetylhydrolase 1b Regulatory Subunit 1 (PAFAH1B1, also known as LIS1), is required for dynein-mediated transport and accumulation of dynein at microtubule plus-ends (Splinter et al., 2012). Finally, Actin Related Protein 10 (ACTR10) has been shown to induce dynein and dynactin interaction (Zhang et al., 2008). Confirmatory results by Western blotting for cytoplasmic dynein 1 light intermediate chain 1 (DYNC1LI1) are shown in Figure S2C. Immunofluorescence microscopy reveals a translocation of DYNC1LI1 to the nuclear membrane upon Nocodazole treatment, which is retained upon LMS extraction (Figure S2D).

### TRIM3: a novel microtubule binding protein

As detailed above, TRIM3, otherwise known as BERP, is clearly identified as an outlier protein, that is de-enriched by Nocodazole treatment. This sensitivity is confirmed by immunoblots showing the corresponding extraction pattern (Figure 3A). It belongs to the TRIM family of E3 ligases, some of which have been previously localised to microtubules (Meroni, 2012). So far, microtubule association has been confined to TRIM proteins that possess a COS motif, FNIII and SPRY/B30.2 domain as their C-terminal domain arrangement (Cox, 2012; Short and Cox, 2006). To determine whether TRIM3 is truly a novel MAP, mouse GFP-TRIM3 was transiently transfected into U2OS cells and co-stained for α-Tubulin, with which it shows a clear co-localisation (Figure 3B). This typical distribution is lost upon Nocodazole treatment but remains evident with Taxol. Thus, we have been able to use our proteomics approach to identify a second novel MAP alongside LGALSL reported above.

**Figure 3.**
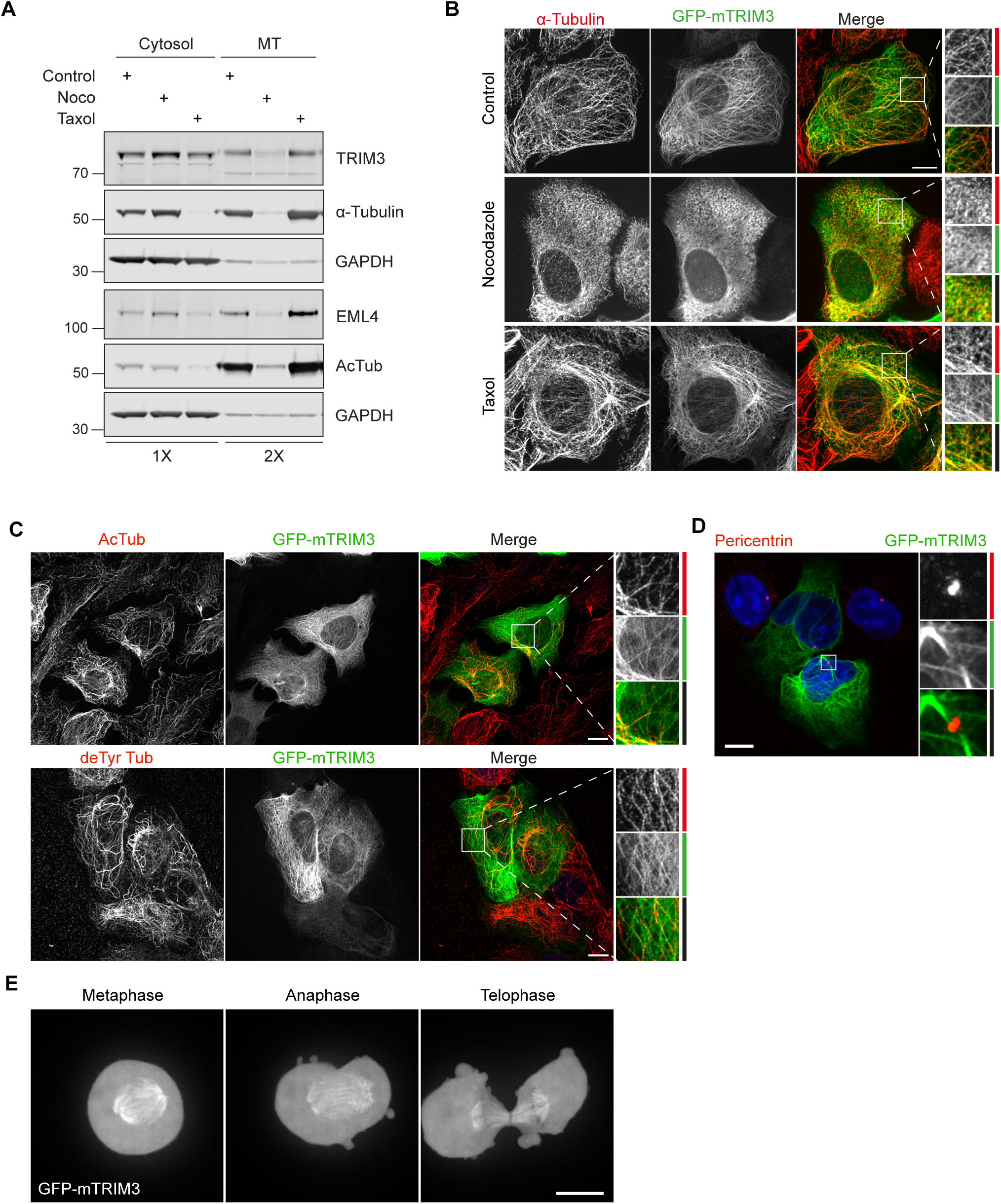
Validation and characterisation of TRIM3 microtubule binding. **A:** U2OS cells were treated with either Nocodazole (6 μM, 1 h) or Taxol (6 μM, 30 min) alongside a DMSO-only treated control sample. Lysis and microtubule stabilisation buffer (LMS) was then added (5 min, 4°C) to remove cytosolic proteins (Cytosol). Residual cellular material, including microtubules, was then collected using 8M urea lysis buffer. Samples were analysed by western blot as indicated, loading twice as much MT-fraction as cytosol. **B:** GFP-mTRIM3 transfected U2OS cells were treated with DMSO (1 h, Control) or Nocodazole (6 µM, 1 h), or Taxol (6 µM, 30 min) before fixation and staining for α-Tubulin (red). **C:** GFP-mTRIM3 transfected cells were fixed and stained for either acetylated Tubulin (AcTub) or detyrosinated Tubulin (deTyr Tub) (red). **D:** GFP-mTRIM3 transfected cells were fixed and stained for Pericentrin (red) and with DAPI (blue). **E:** GFP-mTRIM3 transfected cells were synchronised using thymidine and Nocodazole to arrest cells at prometaphase. Nocodazole was washed out and Z-stacks were acquired every minute (range 15 µm, step 1 µm) as cells progressed through mitosis. A maximum projection is shown. All immunofluorescence experiments were fixed in ice-cold methanol and imaged using a 3i spinning disc confocal microscope, with a Plan-Apochromat 63x/1.4NA Oil Objective M27. Scale bars: 10 µm.

Microtubules are highly decorated with a large number of post-translational modifications (PTMs), and it has come to light that these modifications can play a role in directing which MAPs can associate with which microtubule subsets (Janke and Magiera, 2020). Therefore we analysed whether TRIM3 colocalises with a particular microtubule subset. Figure 3C illustrates that GFP-TRIM3 is not excluded from nor specifically associated with either detyrosinated or acetylated microtubules. However, some of the more intense, seemingly bundled, deTyr microtubule structures are devoid of TRIM3. Furthermore, TRIM3 does not associate with the centrosome (Figure 3D). The microtubule network undergoes major redistribution during mitosis. Following the behaviour of TRIM3 through mitosis in synchronised cells, we observed GFP-TRIM3 on the mitotic spindle from metaphase right through to telophase at which point it equally decorates the central spindle microtubules (Figure 3E, Supplementary Movie 1). The closest paralogue to TRIM3 is TRIM2, and GFP-TRIM2 also shows a clear microtubule association, which has not hitherto been reported (Figure S3A). Like TRIM3, GFP-TRIM2 also associates with a subset of acetylated and deTyr microtubules but is absent from the centrosome (Figure S3B, C).

### The C-terminal region is responsible for TRIM3 localisation to microtubules

TRIM family proteins are defined by their common domains at their N-terminal end and are divided into subfamilies based on their C-terminal domains. Inspection of the 3D structures of TRIM proteins predicted by AlphaFold 2 revealed that all COS-box proteins as well as TRIM3 contain a short α-helix structure shortly after the coiled-coil domain. A schematic and AlphaFold 2 derived structural model of TRIM3 are shown in Figures 4A and B respectively, illustrating the family specific tripartite motif (RING, BB2, Coiled-coil) followed by the short α-helix, a filamin domain and NHL-repeats. Sequence alignments of human TRIM family members reveal that the conserved amino acids within the COS-box, shown to be required for microtubule localisation in other family members, are not present within TRIM3 or TRIM2 (Figure S4 (Short and Cox, 2006)). This suggests that microtubule network localisation is achieved by other means. In order to map which region of TRIM3 is required for microtubule localisation, nine different GFP-tagged truncation or deletion constructs were generated (Figure 4C). Removal of the tripartite motif (ΔRBDC) does not affect microtubule localisation: a truncation retaining the helix, filamin and NHL-repeats in isolation is still able to successfully colocalise with Tubulin (Figure 4C,D). Concordantly the N-terminal Tripartite motif-only construct (ΔH-FNHL) does not bind microtubules and the C-terminal region is therefore solely responsible for microtubule binding. Further analysis of the constructs implicate the C-terminal NHL-repeat region in conjunction with the filamin domain as the minimal determinant of microtubule patterning. Interestingly, the NHL domain alone (ΔRBDCHF) appears to specifically relocate to actin stress fibres.

**Figure 4.**
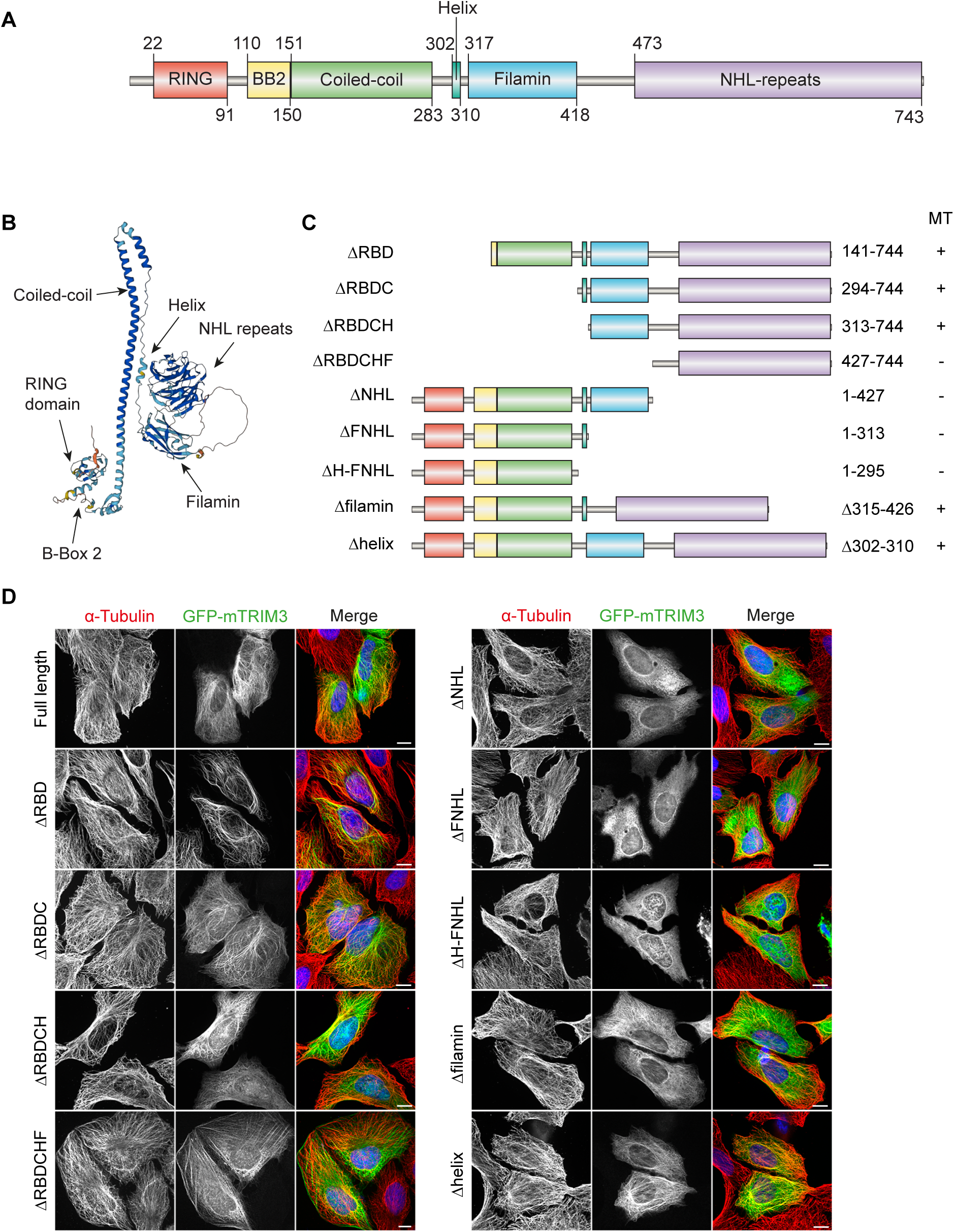
TRIM3 associates with microtubules via its C-terminus. **A:** Schematic representation of full length of TRIM3 and its domains **B:** 3D structural prediction of TRIM3 with domains indicated. Obtained from AlphaFold DB, AF-O75382-F1. **C:** Schematic representation of the TRIM3 deletion constructs used to map the microtubule (MT) localising region. MT binding capability is indicated by +/-. **D:** Representative images of TRIM3 deletion constructs co-stained with α-Tubulin. Images were acquired using a 3i spinning disc confocal microscope with a Plan-Apochromat 63x/1.4NA Oil Objective M27. Scale bars: 10 µm.

### Taxol and Nocodazole treatments reveal an influence of TRIM3 on microtubule properties

We next sought to define a role for TRIM3 in relation to microtubule network properties. TRIM3 was efficiently depleted using siRNA transfection, and immunoblotting and immunofluorescence analyses were performed. Data presented in Figures 5A and B, confirms that levels of total, acetylated or detyrosinated Tubulin are unchanged following depletion, for all indicated time frames. Visual inspection and quantitative analysis does not reveal any obvious perturbation of the microtubule network (Figure 5C-D, Figure S5A). Microtubule networks that are stabilised with acute Taxol treatment undergo an increase in detyrosination and acetylation. To determine whether TRIM3 plays a role in the accumulation rate of these microtubule modifications, TRIM3 was depleted for 72 h before treating with Taxol over a 60 min period. Quantification from three independent repeats reveals that accumulation of detyrosination is not significantly affected, however acetylation is unable to further accumulate under these conditions despite starting from identical baseline levels (Figure 6A,B). Acetylation is a marker of long-lived microtubules, and has previously been shown to be involved in protecting microtubules from mechanical ageing (Portran et al., 2017; Xu et al., 2017). Depletion of the enzyme responsible for microtubule acetylation, Alpha-Tubulin Acetyltransferase 1 (ATAT1) leads to changes in the Nocodazole resistant fraction of microtubules, a finding we have reproduced here (Figure 6C, bottom panel) (Xu et al., 2017). Correspondingly, we find that TRIM3 depletion partially recapitulates this phenotype, wherein residual acetylated microtubules are clearly sparser. This influence of TRIM3 can be accounted for by a 50% reduction in ATAT1 levels upon TRIM3 depletion, which is not due to effects upon transcription as mRNA levels are unchanged (Figure 6D, E). Whilst this provides a plausible mechanistic link for the impact of TRIM3 depletion on microtubule acetylation, how it is effected is currently unclear. ATAT1 is a relatively stable protein (Figure S5B,C) and any direct TRIM3 enzyme-substrate relationship would be expected to result in an increase in ATAT1 levels upon TRIM3 depletion, rather than the observed decrease.

**Figure 5.**
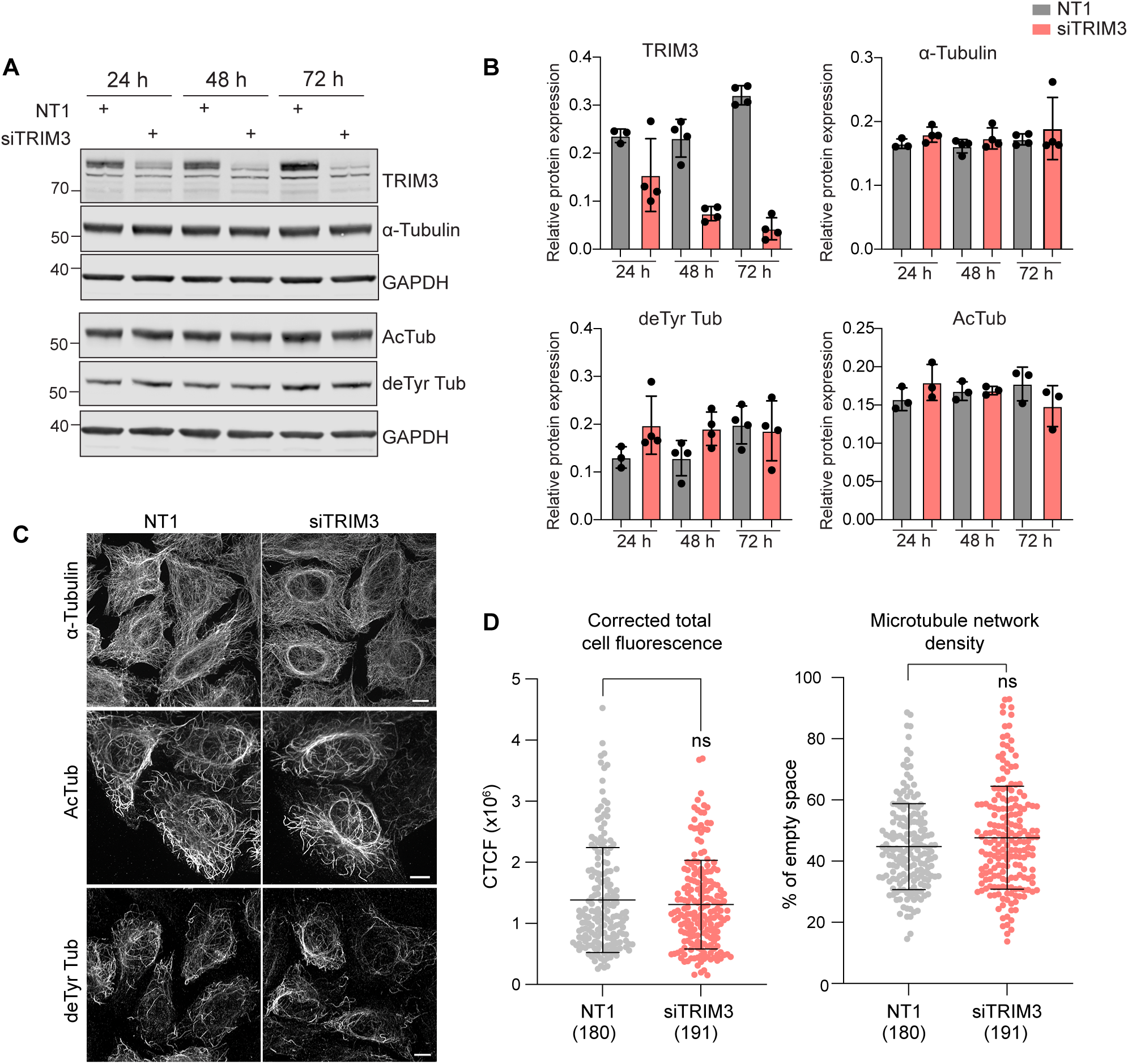
TRIM3 depletion does not affect the global microtubule network. **A:** U2OS cells were transfected with control (NT1) or TRIM3 siRNA (siTRIM3) for 24, 48 or 72 h prior to analysis by western blot. **B:** Quantification of relative expression levels showing mean and standard deviation for ≥3 independent biological experiments for each condition. **C:** U2OS cells were transfected with control (NT1) or TRIM3 targeted siRNA (siTRIM3) for 48 h. Cells were fixed with ice-cold MeOH and stained for α-Tubulin, acetylated Tubulin (AcTub) or detyrosinated Tubulin (deTyr Tub). Images were acquired using a 3i spinning disc confocal microscope with a Plan-Apochromat 63x/1.4NA Oil Objective M27. Scale bar: 10 µm. **D:** Quantitative assessment of the microtubule network showing the corrected total cell fluorescence (CTCF, left) and the percentage of empty space (right) from 3 independent experiments. Mean and standard deviation are shown for the indicated number of cells. Statistical analysis was determined using a student’s t-test.

**Figure 6.**
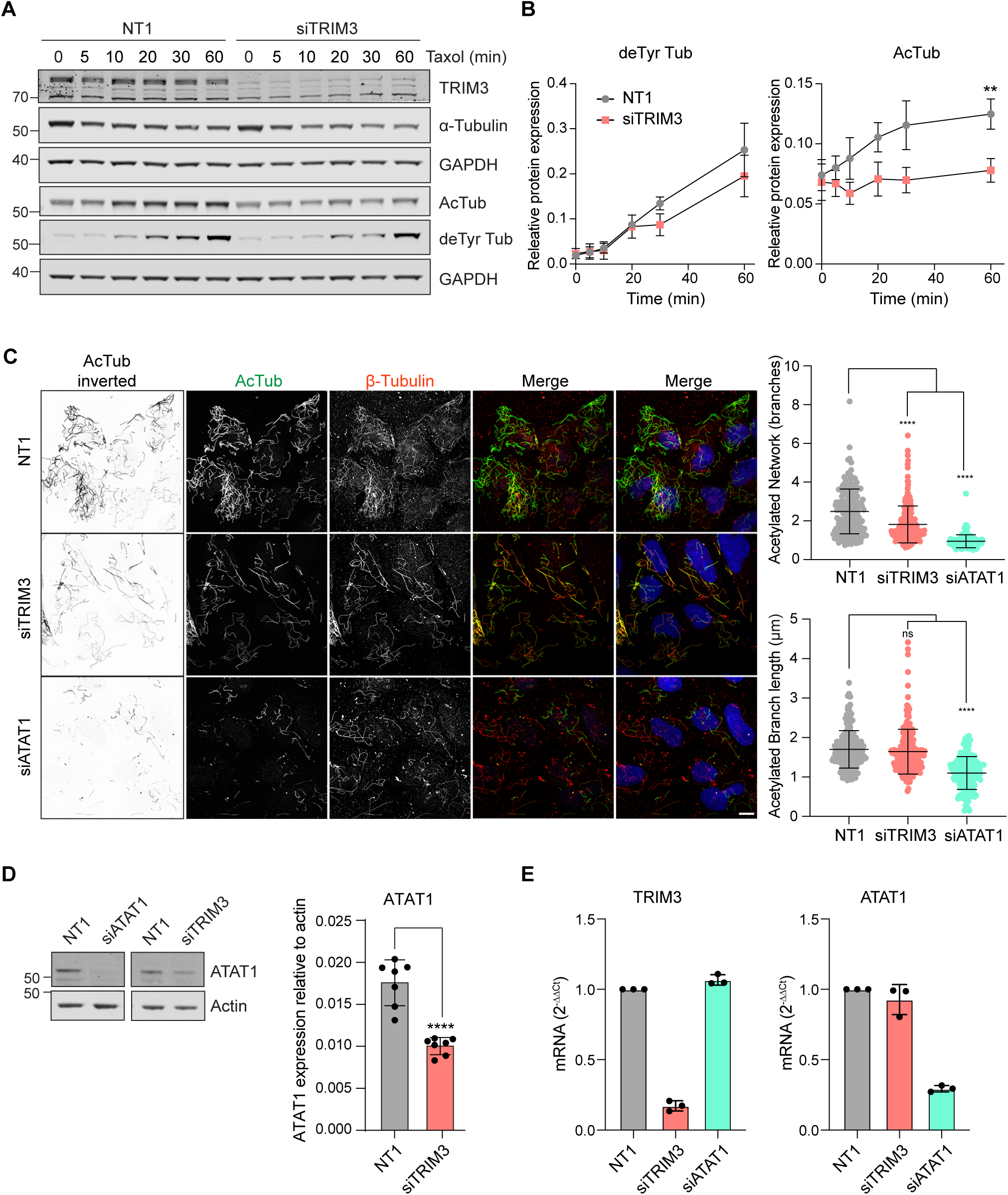
TRIM3 enables drug-sensitive acetylation of microtubules. **A.** U2OS cells were transfected with non-targeting control (NT1) or TRIM3 targeting siRNA (siTRIM3) for 72 h. Transfected cells were treated with 0.5 µg/ml Taxol for indicated timepoints and expression levels of Tubulin and modified Tubulin were analysed. **B.** Quantification of detyrosinated (deTyr) and acetylated Tubulin (AcTub) levels shown in A. Mean and SD of three (deTyr) and four (AcTub) independent experiments are shown. Statistical analysis was determined using 2-way ANOVA with Šídák’s multiple comparisons test. **C:** U2OS cells were transfected with non-targeting control (NT1) or siRNA targeting TRIM3 or ATAT1 for 72 h. Transfected cells were treated with 6 µM Nocodazole for 1 h. Cells were then washed in PHEM buffer to remove the depolymerised microtubules, leaving behind only stable microtubules. Cells were fixed with ice cold MeOH and stained for ß-Tubulin (red) and acetylated Tubulin (AcTub, green). Images were acquired using a 3i spinning disc confocal microscope with a Plan-Apochromat 63x/1.4NA Oil Objective M27. Scale bar: 10 µm. Quantification using MiNA analysis shows the mean number of branches per network (mean branch network) and branch length of acetylated microtubules. Cell numbers analysed: siNT1 (189), siTRIM3 (217), siATAT1 (161). Mean and SD are shown. Significance was determined using a one-way ANOVA with Tukey’s multiple comparison test. **D:** U2OS cells were transfected with control (NT1) or siRNA targeting TRIM3 or ATAT1 for 72 h. Cells were lysed in RIPA buffer and ATAT1 expression levels were analysed by Western blotting. Graph shows quantification of ATAT1 expression levels relative to Actin from 7 independent experiments. Mean and SD are shown. Significance was determined using an unpaired t-test. **E**. U2OS cells were transfected with control or siRNA targeting either TRIM3 of ATAT1 for 72 h. TRIM3 and ATAT1 mRNA levels relative to GAPDH were determined using qRT-PCR. Mean and SD are shown.

## Discussion

Our study provides two novel elements to the analysis of the “MAPome”. Firstly our use of stable isotopes to conduct a triplexed experiment provides a quantitative assessment of enrichment. Secondly, it reflects interactions which occur on microtubule networks in the living cell rather than *in vitro* . However, as with any method that requires a physical separation of bound and unbound, it will be prey to loss of low affinity proteins. Our strategy of differential extraction makes use of the leakage of free proteins versus microtubule associated proteins from the permeabilised cells. This concept has rarely been applied to quantitative proteomic studies, but we recall that differential extraction of membrane proteins by detergents played a key role in the concept of “lipid rafts” and in the identification of caveolin 1 (VIP21) (Kurzchalia et al., 1992).

In our study we are also defining sets of proteins which show the highest sensitivity to two widely used drugs which act upon the microtubule network. This can simply reflect the relative abundance of the microtubule network or changed configuration of Tubulin induced by drug binding. It is the Nocadazole-reduced residual fraction which has been most clearly informative and represents the most novel aspect of this study. It is validated by the high enrichment for microtubule associated proteins amongst the outliers, including most major Tubulin isoforms (Figure 2A,B). It provides a rich resource of uncharacterised proteins from which we have been able to show an association with microtubules for two chosen examples, LGALSL and TRIM3.

Several TRIM proteins have been previously described to localise to microtubules via a signature amino acid sequence, named the COS box, which follows the coiled-coil domain (Short and Cox, 2006). Mutations induced at the beginning and end of this region lead to dissociation from the MT network. TRIM3 does not possess a COS box and we have shown that the MT localising regions lie elsewhere. We identify seven TRIM proteins within our dataset (TRIM3, TRIM24, TRIM56, TRIM28, TRIM33, TRIM23, and TRIM25) of which TRIM3 is the only outlier (Figure S5D). We do not identify TRIM2, the closest relative of TRIM3, with which it is known to heterodimerise (Esposito et al., 2022). However, we were able to show TRIM2 MT association following exogenous expression. The most likely reason we did not find TRIM2 in our data set is its low level of expression in U2OS cells. A high depth quantitative study of the whole U2OS proteome identified 23 TRIM proteins over three orders of magnitude expression levels, but failed to register TRIM2 (Beck et al., 2011).

The TRIM family have been associated with multiple cellular functions with relevance to both cancer and immunity (Hatakeyama, 2011; Ozato et al., 2008). Proposed TRIM3 substrates include Toll Like Receptor 3 and Importin α, (Li et al., 2020; Zhu et al., 2019). It has also been shown to bind to the neuronal Kinesin motor protein KIF21B, whose motility it regulates in a ligase independent manner (Labonte et al., 2013). Our goal in this study was to develop a new approach for gaining a global view of MAPs. Accordingly, we do not provide a complete and detailed mechanistic study of TRIM3 function, but have made some intriguing observations.

Using orthologous assays, we have unveiled a subtle phenotype linking TRIM3 to the accumulation of acetylated microtubules. Our finding that TRIM3 also regulates the levels of the enzyme responsible for acetylation, ATAT1, provides a plausible explanation and starting point for further investigation. We are not proposing ATAT1 as a direct TRIM3 E3 ligase substrate leading to its degradation, as TRIM3 depletion would then result in an increase rather than the observed decrease. Additionally, TRIM3 is a relatively poor E3 ligase in isolation, but can be activated by association with its paralogue TRIM2, which as discussed is not found in our data or in U2OS cells more generally (Esposito et al., 2022). At this point we cannot rule out TRIM3-mediated effects upon ATAT1 translation, particularly as its NHL repeat domain is proposed to confer RNA binding activity as well as the MT-binding shown here (Goyani et al., 2021).

We were surprised to find a coherent set of proteins enriched in the Nocadozole-treated residual fraction. We ascribe this to the Nocadazole-induced relocation of dynein-associated molecules to the nuclear membrane. Such a Nocodazole-induced translocation phenomenon has been previously described for dynein, and reflects conditions in the G2 phase of the cell cycle leading to association of BICD2 and dynein with nuclear pores (Gerlitz et al., 2013; Splinter et al., 2010). This nuclear membrane redistribution has also been well described for another major outlier PAFAH1B1 (also known as LIS1) (Hebbar et al., 2008). In effect Nocodazole may here be mimicking the choreography of mitosis, whereby disassembly of microtubules in prophase is accompanied by dynein-dependent nuclear envelope breakdown (Salina et al., 2002).

Some proteins are at first glance counterintuitively de-enriched by both Nocadazole and Taxol treatments, most notably EB3 (MAPRE3). If a protein is released by Nocodazole treatment why would they also be released upon microtubule stabilisation? In this case, the answer would appear to lie in the specific association of EB proteins with GTP-tubulin at the growing tips of microtubules which are lost upon Taxol stabilisation (Maurer et al., 2011). The dual sensitivity of EB3 in the two drug treated conditions can be rationalised in this way and the assay is reflecting real biological behaviour. Nevertheless, the identification of MT-associated EB3 in control cells may be considered surprising given the high dissociation rate of EB family proteins (Roth et al., 2018). The related protein EB1 shows a similar trend but is below our cut-off value, perhaps reflecting a higher dissociation rate from microtubules in the control condition.

In summary, we have established a new method for identification of MAPs, which we hope will translate into other cell types and prove useful in the investigation of pathological conditions. The ability to culture iPSC neurons at scale will offer an opportunity to transfer our system to a specialised microtubule network (Tian et al., 2019). Our vision is that this method will be one component of a new wave of mass spectrometry based methods to comprehensively detail the MAPome. There is a broad unresolved question as to why multiple TRIM family ubiquitin E3 ligases associate with microtubules, to which we now contribute two new examples. TRIM2 has been previously linked to neuronal polarisation and knock-out mice show evidence of neurodegeneration (Balastik et al., 2008; Khazaei et al., 2011). TRIM2 and TRIM3 are both enriched in the brain and their functions there must be considered in the light of our findings.

## Materials and Methods

### Cell culture

U2OS cells from the European Collection of Authenticated Cell Cultures (ECACC) were grown in Dulbecco’s Modified Eagle Medium (DMEM) supplemented with 10% heat-inactivated foetal bovine serum at 37 °C with 5% CO_2_. Cells were routinely tested for mycoplasma.

### Antibodies

The following primary antibodies were used for western blotting at the indicated concentrations: α-Tubulin (Sigma, T5168, 1:10,000), acetylated α-Tubulin (Sigma, T6793, 1:1000), ATAT1 (Proteintech, 28828-1-AP, 1:500), Detyrosinated α-Tubulin (Abcam, ab32386, 1:1000), DYNC1LI1 (Atlas, HPA035013, 1:1000), EML4 (Cell Signaling Technology, 12156S, 1:1000), GAPDH (Cell Signalling Technology, 2118S, 1:1000). The following primary antibodies were used for immunofluorescence at the indicated concentrations: α-Tubulin (Bio-rad, MCA77G, 1:500), β-Tubulin (Abcam, ab6046, 1:200), acetylated α-Tubulin (Sigma, T6793, 1:1000), Detyrosinated α-Tubulin (Abcam, ab32386, 1:200), DYNC1LI1 (Atlas, HPA035013, 1:1000), MAP4 (Bethyl, A301-488A, 1:1000), Pericentrin (Abcam, ab4448, 1:1000). Secondary antibodies used for western blotting obtained from LICOR Biosciences: Donkey anti-mouse IRDYE 800CW (926-32212), Donkey anti-mouse IRDYE 680CW (926-32222), Donkey anti-rabbit IRDYE 800CW (926-32213), Donkey anti-rabbit IRDYE 680CW (926-32223). Secondary antibodies used for immunofluorescence obtained from Invitrogen: Donkey anti-rabbit AF488 (A21206), Donkey anti-rabbit AF594 (A21207), Donkey anti-mouse AF488 (A21202), Donkey anti-mouse AF594 (A21203), Donkey anti-rat AF594 (A21209).

### Plasmids

Full length LGALSL was amplified by PCR from pBluescriptR-LGALSL purchased from Horizon Discovery (Cambridge, UK) (MHS6278-202809144, clone ID: 5301908). Full length TRIM2, mTRIM3 and mTRIM3 truncations and deletions were amplified by PCR from pcDNA3X(+)MycEGFP-TRIM2 and pcDNA3X(+)MycEGFP-mTRIM3 which were a kind gift from Prof. Germana Meroni (University of Trieste). PCR amplicons were then sub-cloned into BglII and SalI sites of the pEGFP-C1 vector.

### Drug treatments

Nocodazole was used at 330 nM for synchronisation and 6 µM for microtubule depolymerisation. Taxol was used at 6 µM for microtubule stabilisation and 600 nM for accumulation of modifications. Thymidine was used at 2 mM and cycloheximide at 100 µg/ml. The duration of treatments is detailed in respective figure legends.

### Cell lysis, SDS-PAGE and Western blotting

Cells were placed on ice, rinsed twice with ice-cold PBS and lysed in RIPA buffer (150 mM NaCl, 1% Triton-X100, 0.1% SDS, 1% sodium deoxycholate, 10 mM Tris-HCL pH 7.5) supplemented with mammalian protease inhibitors (1:250 v/v) on a rocker on ice for 15 minutes. All samples were suspended in Laemmli sample buffer and boiled (95 °C, 5 minutes). Proteins were resolved on precast NuPAGE Novex 4-12% Bis-Tris Gels in 3-(*N*-morpholino)propanesulfonic acid (MOPS) buffer (*Invitrogen*). Samples were transferred to 0.45 µm nitrocellulose membrane (0.9 A, 1 h) in transfer buffer (0.2 M Glycine, 25 mM Tris, 20% methanol). Membranes were incubated in blocking solution (Tris buffered saline (TBS: 150 mM NaCl, 10 mM Tris, pH 7.5) supplemented with 0.1% Tween-20 (TBST, v/v), and 5% Marvel (w/v) 1 h without and then overnight with primary antibodies, washed 3x for 5 min in TBST before incubating in either IRDye680 or IRDye800-conjugated anti-mouse or anti-rabbit secondary antibodies in blocking solution for 1 h (1:15,000 v/v, *LI-COR*). Visualisation and quantification of Western blots were performed using an Odyssey infrared scanner (LI-COR Biosciences). For western blot quantitation, raw signal values were obtained from image studio software following background subtraction. For quantitation across multiple experiments, raw values were either normalized to the sum of the raw values from each individual blot (Figure 5A and 6B) or normalised to the corresponding Actin bands (Figure 6D).

### Fluorescence microscopy

Cells were seeded onto 22 mm^2^ coverslips in 6-well plates. Cells were fixed with ice cold methanol (MeOH) at -20 °C for 5 min, washed in PBS before incubating with 10% goat serum in PBS for 30 min at room temperature. Primary antibodies (1 h) and secondary antibodies (30 min) were applied sequentially in 5% goat serum, each followed by two 4 min washes with PBS. Coverslips were finally washed in water and mounted onto glass slides using Mowiol supplemented with DAPI. Images were acquired using either a Nikon Ti Eclipse microscope with a CFI Plan Apo 40x objective or a CFI Plan Apochromat VC 60X objective lens, or a 3i Marianas spinning disk confocal microscope (3i Intelligent Imaging innovations, Germany) with a Plan-Apochromat 40x/1.3NA Oil Objective or a Plan-Apochromat 63x/1.4NA Oil Objective M27 or using a Zeiss LSM900 with Airyscan confocal laser scanning microscope using a 63x x 1.4 NA Zeiss Plan Apochromat objective.

### Microtubule extraction protocol

Cells were treated with 6 μM of Nocodazole for 1 h or 6 μM of Taxol for 30 min at 37 °C alongside a DMSO control. The following steps were performed on ice using buffers pre-cooled to 4 °C. Cells were washed in PBS before being incubated in Lysis and Microtubule Stabilisation buffer (LMS: 100 mM PIPES pH 6.9, 5 mM MgCl_2_, 1 mM EGTA, 30% glycerol, 0.1% NP40, 0.1% Triton X-100, 0.1% Tween-20, 0.1% 2-mercaptoethanol (v/v)) supplemented with MPIs (1:250) for 5 min before collection of lysates. The remaining microtubules on the plates were then extracted using 8 M urea, 50 mM Tris, pH 8 (8M urea lysis buffer).

### siRNA transfection

All siRNA treatments were performed the day after cell seeding using Lipofectamine RNAiMAX transfection reagent, in accordance with the manufacturer’s protocol. Cells were incubated for either 48 h or 72 h as specified. Sequences of siRNA oligos used are as follows: ONTARGETplus Non-targeting siRNA oligo#1 (NT1) 5’-TGGTTTACATGTCGACTAA-3’ (Dharmacon, D-001810-01), TRIM3 oligo#1 GUACAGCACAGGCGGCAAA, oligo#2 GCACAUAUGAGCUAGUGUA, oligo#3 GAGCGCCACUGCACACGAA, oligo#4 GAAUGAAAUUGUAGUAACG (Dharmacon, L-006931-00), ATAT1 oligo#1 GUAGCUAGGUCCCGAUAUA, oligo#2 GAGUAUAGCUAGAUCCCUU, oligo#3 GGGAAACUCACCAGAACGA, oligo#4 CUUGUGAGAUUGUCGAGAU.

### Plasmid DNA transfection

All plasmid DNA transfections were performed using 1 µg of plasmid and Genejuice or Lipofectamine 2000 the day prior to collection of experimental data in either a 6-well plate or µ-Dish 35 mm high Ibidi® dishes (Ibidi LCC, Martinsried, Germany) for live cell imaging, with cells at 60-80% confluency at the time of transfection.

### Cell synchronization and live cell imaging

For imaging of TRIM3 on the mitotic spindle, cells were synchronized to increase the mitotic index. Cells were seeded into 35 mm ibidi dishes the day prior to synchronization in prometaphase using 2 mM thymidine (24 h) followed by 330 nM Nocodazole (16 - 18 h). Transfection with plasmid DNA was performed alongside Nocodazole addition. To allow re-entry into mitosis, cells were quickly rinsed three times, then incubated in complete media for 1 hour prior to imaging. Cells in late prometaphase or metaphase were selected and z-stacks were acquired every minute as cells passed through mitosis. Imaging was acquired using a 3i Marianas spinning disk confocal microscope (3i Intelligent Imaging innovations, Germany) with a Plan-Apochromat 63x/1.4NA Oil Objective M27.

### Analysis of stable microtubules

U2OS cells were transfected with siRNA against TRIM3 and ATAT1 alongside a non-targeting control (NT1) for 72 h. Cells were then treated with 6 µM Nocodazole for 1 h to depolymerise the dynamic microtubules whilst maintaining Nocodazole-resistant ones. Cells were washed twice in a buffer containing <u>P</u>IPES, <u>H</u>EPES, <u>E</u>GTA and <u>M</u>gSO_4_ (PHEM: 60 mM PIPES, 25 mM HEPES, 10 mM EGTA and 4 mM Mg SO_4_) before treatment with PHEM buffer containing 0.2% Triton-X 100 for 1 min. This removes the cytosolic Tubulin (depolymerised microtubules) whilst maintaining the intact stable microtubules. Cells were then fixed and stained as described above.

### SILAC labelling

U2OS cells were cultured under standard conditions in DMEM for SILAC supplemented with 10% dialysed FBS and L-proline, Pro0 (200 μg/ml) to prevent conversion of arginine to proline. Differentially labelled amino acids were added to the media to generate SILAC media for 3 conditions: Light (L-lysine, Lys0; L-arginine, Arg0), Medium (L-lysine-^2^H_4_, Lys4; L-arginine-^13^C_6_, Arg6) and Heavy (L-lysine-^13^C_6_-^15^N_2_, Lys8; L-arginine-^13^C_6_-^15^N_4_, Arg10). L-lysine was supplemented at a final concentration of 146 μg/ml whereas L-arginine was supplemented at a final concentration of 84 μg/ml. Cells were cultured for a minimum of 6 passages and analysed by mass spectrometry to confirm amino acid incorporation before experiments proceeded.

### Mass spectrometry

Extracted microtubules from each condition were combined at a 1:1:1 ratio and filtered through 15ml Amicon ultra centrifugal filters (10,000kD MW cut off) at 4000g at RT until the minimum volume was retained. The flow-through was discarded and the sample suspended in sample buffer before being subjected to processing for mass spectrometry. SDS-PAGE was performed until proteins had migrated 1/3 through the gel, then stained with SimplyBlue^TM^SafeStain. The sample lane was cut into 12 equal slices, then into 1 mm cubes and destained using 50 mM Ammonium bicarbonate (Ambic) containing 50% Acetonitrile (ACN, v/v) at 37°C in an Eppendorf ThermoMixer Compact at 900 rpm. Samples were dehydrated (ACN, 5 min) and dried via Speedvac rotary evaporation. Peptides were reduced in 10 mM dithiothreitol (56 °C, 1 h), alkylated in 55 mM iodoacetamide (RT, 30 min), washed in 100mM Ambic (15 min) then repeated in 50 mM Ambic with 50% ACN. Samples were dehydrated in ACN, followed by Speedvac evaporation. Peptides were cleaved with 3 μg trypsin per sample lane in 40 mM Ambic and 9% ACN (37 °C, overnight) then extracted using ACN (30°C, 30 min), followed by 1% formic acid (FA room temperature, 20 min) and then ACN again. Supernatants were dried in a Speedvac overnight and stored at -20°C until analysis.

Samples were desalted using C18 tips (Rappsilber et al., 2003). Briefly, samples were resuspended in 50 µl 0.1% FA. C18 tips were first activated with Methanol, equilibrated twice with 0.1% FA, then samples were loaded, and peptides were washed with 0.1% FA. Samples were then eluted with 80% ACN / 0.1% FA and dried in the speed vac. Finally, they were reconstituted in 0.1% FA. Peptides were then fractionated on a QExactive HF plus column coupled to a nanoHPLC Ultimate 3000 using a pre-concentration onto a nano-column configuration. An Acclaim PepMap 100 (75 µm, 2 cm) was used to do the pre-concentration and an Acclaim PepMap RSLC (Rapid separation liquid chromatography, 75 µm, 15 cm) was used for peptide separation. Total run time was 118 min with a gradient from 4% to 40% Buffer B [Buffer B: 80% ACN / 0.1% FA], in the following steps: 2% for 8 min, 2% to 40% in 80 min, 90% for 10 min and an equilibration step with 2% for 20 min. The MS was operated in a data dependent manner using a loop count of 12. MS1 was acquired in a scan range from 375-1500 m/z, with a resolution of 120000, an AGC target of 3x10^6^ and a maximum IT (injection time) of 120 ms. MS^2^ was acquired at a resolution of 17,500, an AGC target of 1x10^5^ and maximum IT time of 60 ms. Dynamic exclusion was set to 20 s and ion with charge states of +1 and greater than +6 were excluded. Searches were performed using MaxQuant, version 1.6.17.0, against the human database. Carbamidomethylation of cysteines (+57 Da) was used as a fixed modification and oxidation of Methionine (+16 Da) and Acetylation of the N-terminal (+42 Da) as variable modifications. Multiplicity was set to 3 and the labels were as follow: Medium labels - Arg6;Lys4, Heavy labels - Arg10;Lys8.

### Microtubule analysis

The MiNa (Mitochondrial Network Analysis) plugin for Fiji (Valente et al., 2017) was utilised to characterise the interconnectivity of the microtubule network. The corrected total cell fluorescence (CTCF) of the microtubule network was analysed using Fiji and calculated using the following equation: CTCF = integrated density – (area of selected cell x mean fluorescence background). The microtubule network density was analysed using Fiji to calculate the percentage of empty space by applying a threshold value to the images based on the control sample.

### Statistical analysis

For western blot quantifications, band intensities were measured using Image Studio Software. All statistical analysis was carried out using GraphPad Prism. Error bars: mean ±SD. ns>0.05, *p≤0.05, **p≤0.01, ***p≤0.001, ****p≤0.0001).

## Acknowledgements

HG and JGN are recipients of PhD studentships from the Wellcome Trust and North West Cancer Research respectively. MC is a recipient of a Royal Society Industry Fellowship INF\R2\212031. We thank the Liverpool Centre for Cell Imaging for enabling this work.

